# Machine Learning-assisted Raman Spectral Analysis of Serotonin-responsive ssDNA-SWCNT Nanosensor for Improved Selectivity against Dopamine

**DOI:** 10.1101/2025.10.02.679925

**Authors:** Yunseo Choe, Sanghwa Jeong

## Abstract

Serotonin (5-hydroxytryptamine, 5-HT) plays critical roles in neuromodulation, yet current detection methods struggle for real-time sensing of 5HT with high sensitivity and selectivity. We previously developed nIRHT (near-infrared serotonin nanosensor), which consists of ssDNA-wrapped single walled carbon nanotube that sensitively detects 5-HT. However, nIRHT’s fluorescence response cannot discriminate between 5HT and dopamine (DA), limiting its practical applications. In this study, Raman spectroscopy combined with machine learning overcomes this selectivity challenge. G-band spectral features revealed distinct signatures for 5HT versus DA binding to nIRHT, with DA causing greater G^-^ band suppression. We employed differential Raman (ΔRaman) to isolate analyte-specific spectral changes and trained three machine learning models for classification. The random forest model with ΔRaman achieved optimal performance with 95.8% accuracy, significantly outperforming models using raw Raman spectra. This approach showed improved specificity, with negligible responses to acetylcholine, GABA, and glutamate, and achieved a detection limit of 0.1 μM suitable for physiological applications. This Raman based approach transforms the non-selective nIRHT fluorescence sensor into a platform capable of robust neurotransmitter discrimination, overcoming selectivity issues in SWCNT-based molecular sensing.

## 1. Introduction

Serotonin (5-hydroxytryptamine, 5-HT) is a critical neurotransmitter regulating mood, cognition, sleep, and appetite, with dysfunction implicated in depression, anxiety, and other neuropsychiatric disorders^1^. Real-time monitoring of serotonin dynamics in biological systems remains challenging due to the millisecond timescale of synaptic transmission and the complex chemical environment of neural tissue^2^. Despite significant advances in neurotransmitter detection technologies, achieving both high sensitivity and molecular selectivity for serotonin continues to pose substantial analytical challenges. Current serotonin detection methods span from traditional analytical techniques to emerging nanosensor platforms. High-performance liquid chromatography with electrochemical detection (HPLC-ED) remains the gold standard, achieving detection limits of 0.056 nM with excellent reproducibility^3^. However, HPLC requires extensive sample preparation and 15-30 minute analysis times, precluding real-time monitoring□. Fast-scan cyclic voltammetry (FSCV) enables subsecond temporal resolution at carbon fiber microelectrodes but suffers from electrode fouling and limited selectivity between structurally similar neurotransmitters□,□. Recent developments in genetically-encoded fluorescent sensors, including iSeroSnFR□ and GRAB-5HT□, achieve remarkable sensitivity and dynamic range but require genetic modification of target tissues, limiting clinical translation.

Single-walled carbon nanotube (SWCNT) optical sensors have emerged as promising platforms for label-free neurotransmitter detection□. DNA-functionalized SWCNTs exhibit near-infrared (nIR) fluorescence modulation upon analyte binding, enabling detection in the tissue-transparent window (900-1400 nm)^1^□,^11^. Landry group developed the nIRHT sensor through systematic evolution of ligands by exponential enrichment (SELEC), screening ∼10^1^□ unique ssDNA sequences to identify serotonin-responsive constructs^12^. This sensor achieves micromolar sensitivity with 200% fluorescence enhancement upon serotonin binding. Similarly, Strano group pioneered corona phase molecular recognition (CoPhMoRe), demonstrating that (GT)□□-wrapped SWCNTs detect dopamine at 11 nM concentrations^13^. Our group recently advanced this field through machine learning-guided optimization, achieving 2.5-fold sensitivity improvements using high-throughput screening of systematically mutated sequences^1^□.

Despite these advances, a fundamental limitation persists: insufficient selectivity between serotonin and dopamine. Both neurotransmitters contain aromatic rings and ethylamine chains, differing by only a single hydroxyl group^1^□. Current fluorescence-based SWCNT sensors typically achieve selectivity ratios below 5:1, inadequate for distinguishing co-released neurotransmitters in synaptic environments^1^□. This selectivity challenge is particularly critical given that serotonin and dopamine often colocalize in brain regions governing reward and motivation^1^□.

Raman spectroscopy offers complementary chemical information through vibrational mode analysis of the SWCNT-analyte complex^1^□. The G-band (∼1580 cm□^1^) of SWCNTs splits into G□ and G□ components, with frequencies sensitive to molecular adsorption and electronic doping^1^□. Previous studies demonstrated that non-covalent surfactants on SWCNT induce distinct Raman spectral changes in G-band region^20^.

In this work, we demonstrate machine learning-assisted Raman spectral analysis of the nIRHT sensor to achieve enhanced selectivity between serotonin and dopamine. We systematically characterized concentration-dependent Raman responses from 0.01 to 100 μM, revealing distinct spectral fingerprints for each neurotransmitter. Three machine learning models—convolutional neural network (CNN), support vector machine (SVM), and random forest (RF)—were trained on spectral datasets to classify analyte identity. Critically, we introduce differential Raman processing, which removes baseline signals to isolate analyte-specific features. The optimized RF model with DR processing achieves 95.8% classification accuracy, representing a significant advance in molecular discrimination. This multimodal approach combining nIR fluorescence sensitivity with Raman chemical specificity provides a pathway toward real-time, selective serotonin monitoring in complex biological environments.

## 2. Result and Discussion

### 2.1. Raman Spectroscopic Analysis Reveals Distinct Signatures for Serotonin and Dopamine

The E6#9 ssDNA-functionalized SWCNT nanosensor (nIRHT), previously developed through SELEC methodology, demonstrates reversible nIR fluorescence enhancement upon 5HT binding, enabling real-time 5HT imaging in vitro and ex vivo^12^. The E6#9 sequence (5’-CCCCCCAGCACCAGACAGCACACTCCCCCC-3’) wraps around SWCNTs to create binding sites for 5HT. However, this sensor lacks specificity: both 5HT and DA results in comparable intensity increase (Fig. 1B), limiting its utility to discriminate between DA and 5HT.

**Figure 1.**
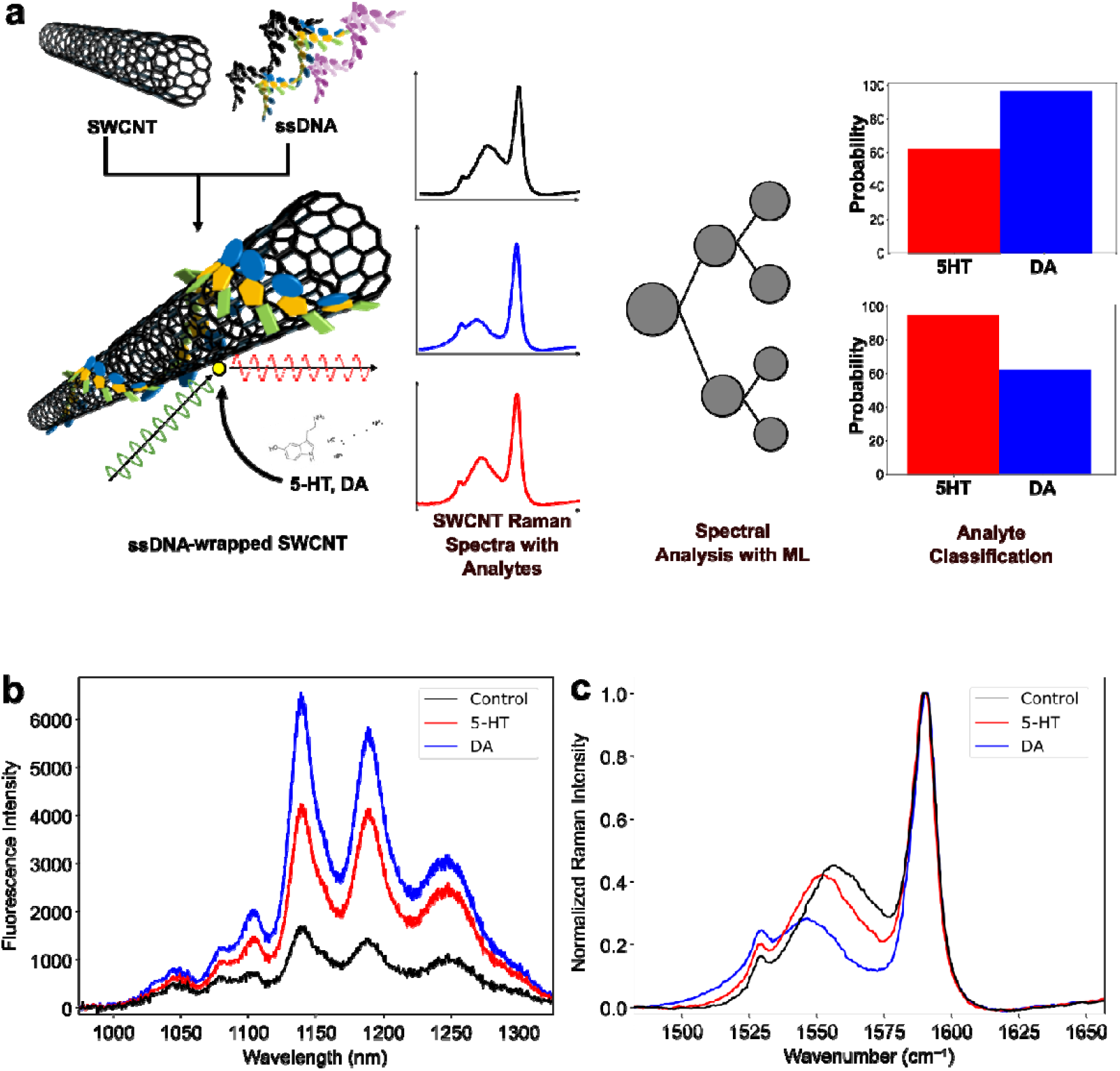
(a) Scheme for the classification of serotonin (5-hydroxytrypamine, 5HT) and dopamine (DA) by using machine learning-assisted Raman spectral analysis on near-infrared 5HT-responsive ssDNA-SWCNT (nIRHT) nanosensor (b) Fluorescence spectra of nIRHT solution before (black) and after incubation with 100 μM 5HT (blue) and DA (red). (c) Normalized Raman G band spectra of nIRHT solution before (black) and after incubation with 100 μM 5HT (red) and DA (blue).

To overcome this limitation, we employed Raman spectroscopy to investigate molecular-level interactions between neurochemicals and nIRHT nanosensors. Raman analysis of nIRHT nanosensor solution revealed distinct spectral features in the G-band region (∼1580 cm^-1^, which originates from in-plane carbon vibrations in the graphene-like structure of SWCNTs (Fig. 1C). The G band could be splitted into two components: G^+^ band (1580 cm^-1^) corresponds to longitudinal optical phonons along the nanotube axis, and G^-^ band (1540-1570 cm^-1^) corresponds to circumferential vibrations^21, 22^. Previous studies established that covalent modifications such as oxidation shift G-band frequencies through disruption of the sp^2^ carbon network^23^. We observed that non-covalent 5HT and DA adsorption onto nIRHT modulates G-band features with different modes. DA causes pronounced suppression of the G□ band compared to 5HT, providing a spectroscopic fingerprint for neurotransmitter discrimination. DA likely forms stronger π-π interactions with the SWCNT surface due to its catechol moiety, inducing greater electronic perturbation manifested as enhanced G□ suppression. In contrast, 5HT’s indole ring system produces weaker electronic coupling, preserving more of the intrinsic Raman signature.

Raman spectral changes across seven concentrations were measured in the range of 0.01, 0.1, 1, 10, 100 μM, encompassing physiological neurotransmitter levels (Fig. S1). DA exhibited pronounced concentration-dependent G□ band suppression, with maximum attenuation at 100 μM that progressively diminished at lower concentrations (Fig. 2A). 5HT induced qualitatively similar but quantitatively weaker concentration-dependent changes, maintaining distinguishable spectral features even at low concentrations (Fig. 2B).

**Figure 2.**
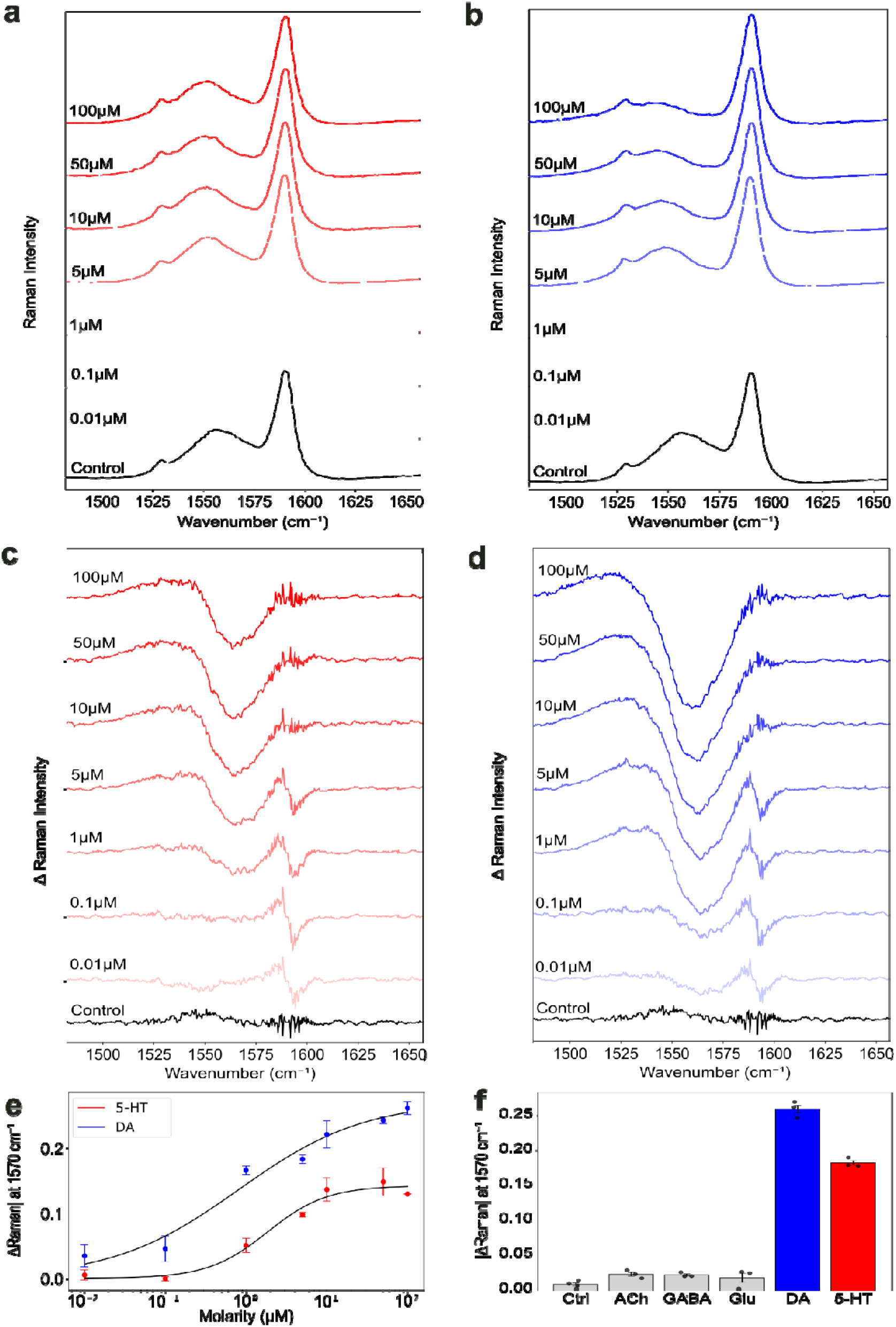
(a) Raman spectra after incubation with 5HT at different concentrations of 100, 50, 10, 5, 1, 0.1 and 0.01 μM. (b) Raman spectra after incubation with DA at different concentrations of 100, 50, 10, 5, 1, 0.1 and 0.01μM. (c) Differential Raman (ΔRaman) spectrum of 5HT at seven concentrations, each 100, 50, 10, 5, 1, 0.1 and 0.01 μM. The lighter the color, the lower the concentration. (d) ΔRaman spectrum of DA at seven concentrations, each 100, 50, 10, 5, 1, 0.1 and 0.01μM. (e) Concentration-dependent ΔRaman at 1570 cm^-1^ for 5HT and DA. Data is fitted with the Hill equation. (f) ΔRaman intensity value at 1570 cm^-1^ for different neurotransmitters at 10 μM, which was treated with ACh, GABA, Glu, DA, 5HT, and 1X PBS only (Ctrl).

For quantification of analyte-specific spectral changes, we calculated differential Raman (ΔRaman) spectra by subtracting baseline measurements from analyte-exposed samples.

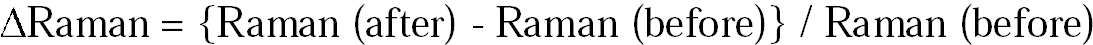

At 100 μM, DA and 5HT generated distinct ΔRaman peak patterns (Fig. 2C and 2D), confirming that each neurotransmitter induces unique spectroscopic shifts despite similar fluorescence responses. These differential signatures arise from varying molecular orientations and binding geometries on the SWCNT surface.

ΔRaman spectra at 1570 cm^-1^ yielded a concentration-dependent curve fitted to the Hill equation (Fig. 2E). The calculated dissociation constants (K_d_) were 2.06 μM for 5HT and 0.89 μM for DA. Both neurochemicals showed sigmoidal dose-response curves with signal saturation above 100 μM. The distinct spectral patterns will enable reliable classification between 5HT and DA, which would be difficult in fluorescence-based detection where both analytes produce indistinguishable responses. The temporal response of nanosensors were measured against 5HT and DA (Fig. S2). The response time is below 1 s for both analytes, which showed the rapid recognition of neurochemicals on SWCNT corona phase.

To evaluate the selectivity of nIRHT’s Raman response to other neurochemicals, we examined spectral changes induced by other major neurotransmitters: acetylcholine (ACh), γ-aminobutyric acid (GABA), and glutamate (Glu). While previous fluorescence studies showed minimal responses to these analytes, their effects on Raman spectra had not been characterized. Relative ΔRaman were calculated relative to baseline measurements (Fig. 2F). ACh, GABA, and Glu produced negligible Raman spectral changes, with ΔRaman values remaining near zero across the entire spectral range. This absence of spectral modulation contrasts sharply with the pronounced G□ band suppression observed for 5HT and DA. This orthogonal validation confirms that the Raman spectral changes are specific to catecholamine and indoleamine neurotransmitters rather than general ionic effects.

These results demonstrate that Raman spectroscopy provides orthogonal chemical information complementary to fluorescence measurements, transforming the nIRHT sensor from a sensitive but non-selective detector into a platform capable of molecular discrimination. The concentration-dependent spectral evolution and analyte-specific ΔRaman signatures establish the foundation for machine learning-based classification algorithms to achieve robust neurotransmitter identification in complex biological environments.

### 2.2. Machine-learning-assisted Classification between Serotonin and Dopamine from Raman Spectral Data

The differential Raman spectra were theoretically distinct for 5HT and DA, while it is challenging to discriminate visually at physiological concentrations. For more robust classification, we implemented machine learning algorithms including convolutional neural network (CNN), support vector machine (SVM), and random forest (RF) to analyze the spectral datasets (Fig. 3A). The training dataset consists of 208 experimental Raman spectra collected across seven concentrations (0.01-100 μM): 76 5HT response, 77 DA response, and 55 control measurements treated with PBS buffer only. Due to the limited dataset size, we employed 5-fold cross-validation to maximize training efficiency and minimize overfitting. The dataset was split into training, validation, and test sets in a 6:2:2 ratio. Performance metrics were calculated as the mean across all validation folds to ensure statistical robustness. Initial evaluation compared classification accuracy (ACC) across the three models using both normalized Raman spectra and ΔRaman as explained in the previous section (Fig. 3B). For normalized Raman spectra, RF achieved the highest ACC at 87.9%, while SVM showed the lowest performance at 71.6%. ΔRaman processing significantly enhanced model performance across all algorithms. The RF model maintained superior performance with 95.8% accuracy, while CNN and SVM showed significant improvements of 9% and 14% rather than the normalized Raman model, respectively. This enhancement demonstrates that baseline subtraction effectively removes systematic noise and helps to focus analyte-specific spectral features for classification. Based on these results, we selected the ΔRaman-based RF model as our optimal classifier model.

**Figure 3.**
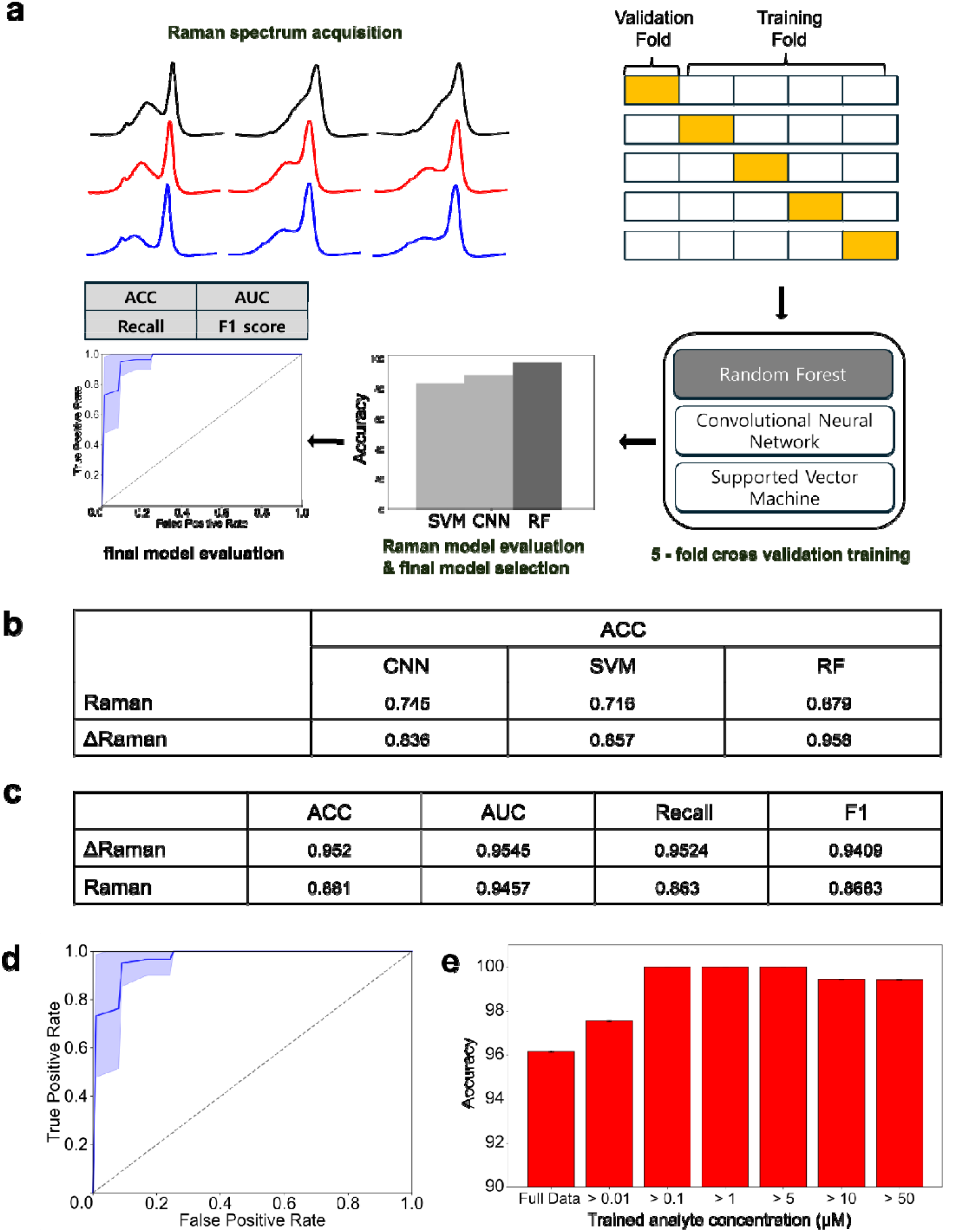
(a) Scheme of machine learning analysis of nanosensor Raman spectral data. After acquisition of spectrum, Raman spectra were trained by three models of CNN, SVM, and RF, in 5-fold cross validation, and evaluated by ACC to select the best model of them. After selection of the best model, the model was assessed by general performance metrics. (b) Accuracy (ACC) performance for each data type and model. (c) General performance metrics at ΔRaman-based RF model and pristine Raman-based RF model. (d) Receiver Operating Characteristic (ROC) curve of ΔRaman-based RF model. (e) ACC of ΔRaman-based RF model depending on different training dataset. Full data uses all concentration data, and >X μM implies to include the truncated dataset which possesses the dataset using more than X μM concentration of 5HT and DA.

We also evaluated normalized Raman and ΔRaman RF models using standard performance metrics including ACC, area under the curve (AUC), recall and F1 score (Fig. 3B). The normalized Raman RF model achieved 94.6% AUC, 88.1% ACC, 86.8% F1 score, and 86.3% recall. However, the ΔRaman-based model showed superior and more balanced performance across all metrics with 95.5% AUC, 95% ACC, 94.1% F1 score, and 95.2% recall. The receiver operating characteristic (ROC) curve for ΔRaman RF model showed excellent discrimination capability (Fig. 3D)

To establish the practical detection range, we evaluated model performance as a function of analyte concentration (Fig. 3E). We evaluated classification performance by sequentially excluding training dataset at each concentration to determine the limit of detection (LOD) indirectly. In comparison to 94.1% ACC without any exclusion, ACC increased to 97.10% after excluding 0.01 μM data. It implied Raman spectral change by 0.01 μM 5HT is somewhat indistinguishable from baseline measurements. While the model trained with 0.1 μM and above, model ACC was saturated between 97-100%, establishing 0.1 μM as the practical LOD for reliable classification. This detection limit aligns with physiological 5HT concentration in synaptic environments^24^, demonstrating the practical applicability of our approach for biological applications.

These results establish that machine learning-assisted Raman analysis will improve the nIRHT sensor from a sensitive but less-selective detector into a platform enabling robust neurotransmitter discrimination. The combination of ΔRaman processing and RF classification provides a robust analytical framework that overcomes the fundamental limitations of fluorescence-based approaches.

## 3. Conclusion

We successfully demonstrated that machine learning-assisted Raman spectroscopy enables the nIRHT nanosensor to discern 5HT from DA with 95.8% accuracy. This approach addresses the fundamental selectivity limitation that has constrained SWCNT-based neurotransmitter sensing. While the nIRHT nanosensor exhibits decent sensitivity through near-infrared fluorescence, it cannot fully distinguish between 5HT and catecholamines. By incorporating Raman spectroscopy as an orthogonal detection modality, we can enhance the molecular specificity necessary for practical applications. The negligible Raman responses to Ach, GABA, and Glu further strengthen the sensor specificity.

Future development should focus on integrating fluorescence and Raman detection in a single platform for simultaneous sensitivity and selectivity, developing deep learning architectures for enhanced classification of neurotransmitter mixtures. This multimodal sensing paradigm combining SWCNT nanosensor, fluorescence imaging, and Raman spectroscopy provides a robust framework for addressing selectivity challenges in molecular sensing and opens new possibilities for real-time monitoring of neurotransmission dynamics in biological systems.

## 4. Methods

### 4.1. Fabrication of ssDNA-functionalized SWCNT nIRHT Nanosensors

The nIRHT solution was synthesized using tip-sonication of SWCNT and ssDNA in 1X PBS buffer. At first, we mixed 1 mg SWCNT (HiPCo raw, Nanointegris) and 100 μl of 1 mM E6#9 ssDNA (sequence = CCCCCCAGCACCAGACAGCACACTCCCCCC) in a 1X PBS 900 μl. Then, this solution was sonicated by a bath sonicator at 5 min. Sequentially, this mixture was tip-sonicated at 50% amplitude for 30 min in an ice bath (3 mm probe, VCX130, SONICS). After sonication, the solution was centrifugated at 21500 rcf for 1 h to precipitate non-dispersed SWCNT. A supernatant of 850 μl was collected. The concentration of the SWCNT suspension was calculated by measuring its absorbance at 632 nm with an extinction coefficient for SWCNT of 0.036 (mg/L)^−1^ cm^−1^. After measuring the concentration, the nIRHT solution sensor was stored in 4 □.

### 4.2. Raman Spectra Measurement

Raman spectra of nIRHT dispersion were collected by RAMANtouch (Nanophoton) at 532 nm laser excitation. The samples were loaded into a 96-well plate before the Raman spectrum acquisition. The spectra of nIRHT dispersion (90 μL) in the PBS buffer were measured as the baseline. Then, each sensor solution was spiked with 10 μL of analyte solution for the final concentration of 0.01, 0.1, 1, 5, 10, 50 and 100 μM. The Raman spectra were measured after 5 min of analyte addition. For each concentration, samples were prepared in triplicate.

### 4.3. Machine Learning-based Data Analysis

To classify the Raman response to 5HT and DA, we applied three machine learning algorithms using Python with Numpy, pandas, and TensorFlow packages. The total dataset was composed of 208 Raman spectra, which contained 77 DA response, 76 5HT response, and 47 control group which was treated by 1X PBS only. Raw Raman spectra in the G-band region underwent sequential pre-processing steps: background subtraction, normalization at G^+^ band peak, and noise-reduction by median filter. Differential Raman (ΔRaman) spectra were calculated by subtracting the normalized control spectrum from each analyte-treated spectrum to isolate spectral changes.

Three classification models were developed for ternary classification (5HT, DA, or control): Convolution Neural Network (CNN), Random Forest (RF), and Supported Vector Machine (SVM). The dataset was split into training, test, and validation set in a 6:2:2 ratio. Due to the small size of data, 5-fold cross validation was employed. CNN was composed of three convolution layers and two fully connected layers. Each convolutional layer used 1D convolution, and had a ReLU layer as activation function and max pooling layer, whose pooling size was 2. RF and SVM were trained using default hyperparameters. Especially, SVM model employed a radial basis function (RBF) kernel for non-linear classification.

Each model was trained on five different fold combinations, with performance metrics averaged across all folds to ensure statistical robustness. The model with highest accuracy on validation data was selected for comprehensive analysis. Final performance was evaluated on the test set to verify general capability.

To determine the limit of detection indirectly, we performed concentration-dependent analysis by sequentially excluding lower concentration data from the training set. The model was trained on data from higher concentration than criteria (>X μM) and tested on all concentrations including those below criteria.

## Acknowledgement

This work was supported by a 2-Year Research Grant from Pusan National University.

