## Supplementary material for "Machine Learning-assisted Raman Spectral Analysis of Serotonin-responsive ssDNA-SWCNT Nanosensor for Improved Selectivity against Dopamine": SI figure

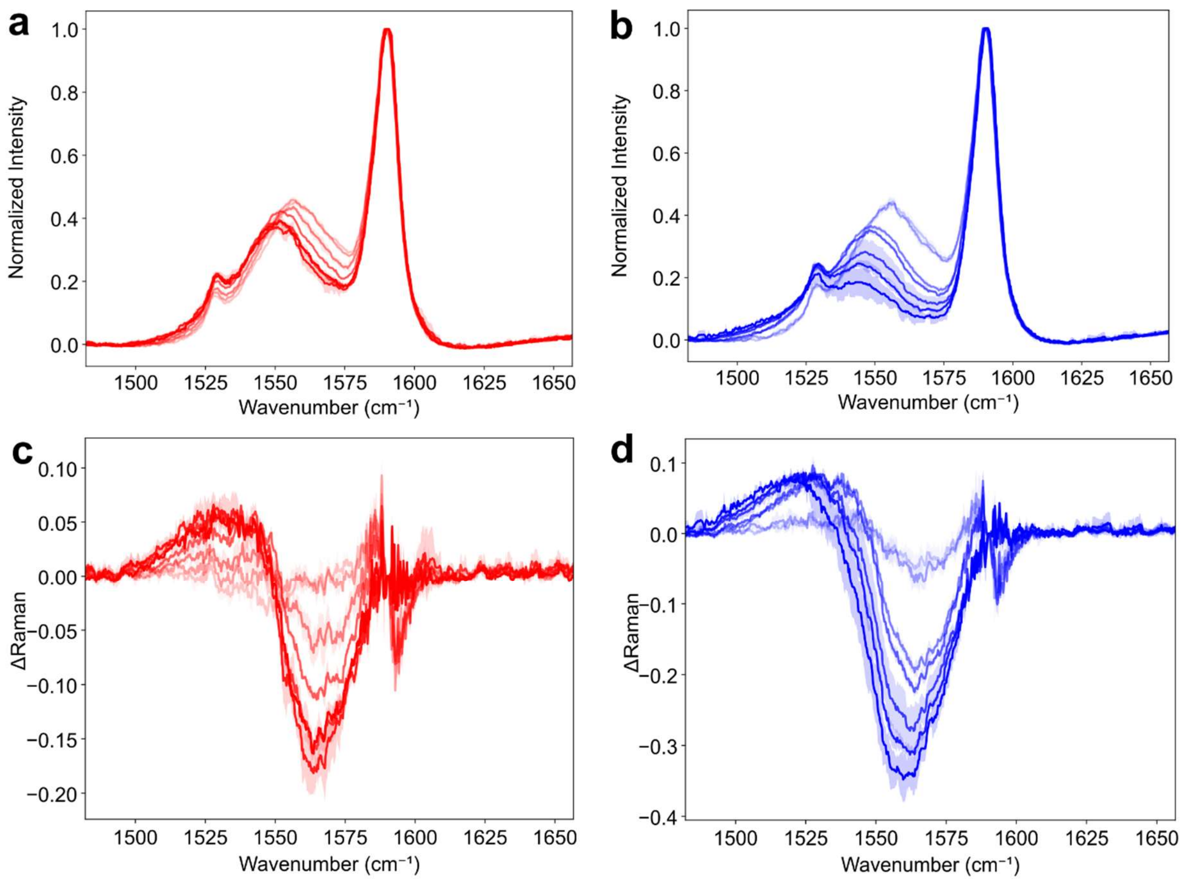


Figure S1. (a) Raman spectra after incubation with 5HT at different concentrations of 100, 50, 10, 5, 1, 0.1 and 0.01 μM. (b) Raman spectra after incubation with DA at different concentrations of 100, 50, 10, 5, 1, 0.1 and 0.01μM. (c) Differential Raman spectrum of 5HT at seven concentrations, each 100, 50, 10, 5, 1, 0.1 and 0.01 μM. (d) Differential Raman spectrum of DA at seven concentrations, each 100, 50, 10, 5, 1, 0.1 and 0.01μM. The lighter the color, the lower the concentration.

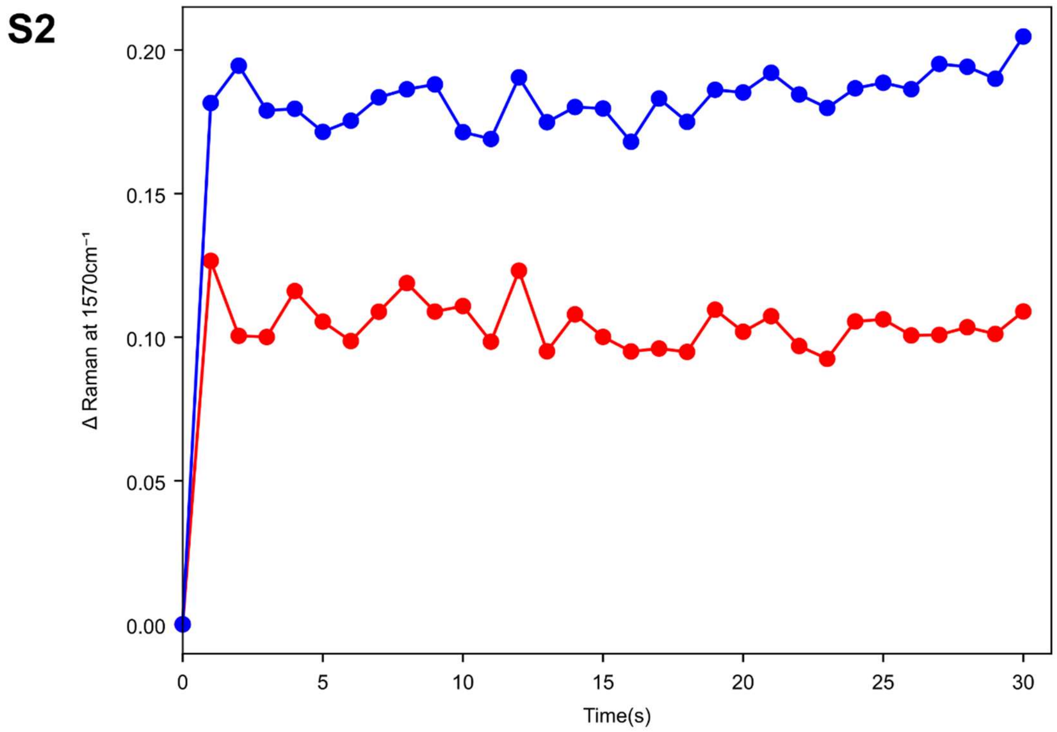


Figure S2. Time-dependent change of differential Raman signal after 5HT (red) and DA (blue) addition.
